# Millet-based supplement restored microbiota diversity of acute undernourished pigs

**DOI:** 10.1101/2019.12.13.875013

**Authors:** Xuejing Li, Yan Hui, JunLi Ren, Yanni Song, Songling Liu, Lianqiang Che, Xi Peng, Xiaoshuang Dai

**Author notes:** Contributing equally to this work (Xuejing Li, Yan Hui).

## Abstract

The strong connection between undernutrition and gut microbiota (GM) has enabled microbiota-targeting to become an evolving strategy, which witnessed urgent need for fortified formula of supplementary food in undernutrition therapy. Using undernourished pigs as models, we investigated how corn- and millet-based nutritional supplement acted differently in modulating microbiota. Undernourished pigs at age of 9 weeks were fed with pure maize diet (Maize), corn-based (CSB+) and millet-based (MSB+) supplementary food for 3 weeks. Compared with Maize group, both CSB+ and MSB+ improved serum total protein and globulin level, but no physiological improvement was observed by short-term food intervention. MSB+ shown superior influence on GM immaturity which were more normally-grown like at both structural and functional level. Higher level of Bacteriodetes, Firmicutes and lower level of Preteobacteria were detected in MSB+ fed piglets in contrast with CSB+. *Lachnospira.spp* was significantly raised after nutritional intervention, indicating high correlation with the undernutrition-associated phenotype. Thus, especially from the GM aspect, millet could be one promising source to help undernourished children reconstruct balanced microbiota in short therapeutic term.

## Introduction

Undernutrition accounts for nearly half of the deaths (∼ 3.1 million) among children under 5 years old in low- and middle-income countries [1]. In 2017, 7.5% of children under age of 5 were affected by wasting syndrome, with regional prevalence ranging from 1.3% in Latin America to 9.7% in Asia[2]. Moreover, more than 90% of global children stunting occurred in Africa and Asia2. Undernutrition usually develops from inadequate protein and micronutrients intake due to unavailability of food [3,4]. Epigenetic modification during pre- and neo-natal period, such as metabolic imprinting, will also increase the risk of malnutrition and diet-related non-communicable diseases later in life [5,6].

CSB+ is one widely-applied ready-to-use supplementary food (RUSF) to treat undernourished children. But fortified therapeutic food is still needed as the undernourished children still have anemia or concurrency of undernutrition in a short period after CSB+ treatment [7,8]. In the last years, increasing evidences accumulated to draw more attention toward the role of the gut microbiota in the development of malnutrition. The gut microbiota works as a virtual organ notably involved in the regulation of several host parameters, including polysaccharide digestion, immune system development [9], defense against infections, synthesis of vitamins and fat storage, etc [10,11]. More recently, Blanton et al emphasized clearly that there existed a causal relationship between immaturity of the gut microbiota and undernutrition. Two kinds of invasive species, Ruminococcus gnavus and Clostridium symbiosum transferred form undernourished donors, could ameliorate growth and metabolic abnormalities in recipient animals [12].

Millet is one of the most important drought-resistant whole grain in arid and semiarid areas of Asia and Africa. It has attracted specific attention on account of the excellent nutritional value and potential health benefits such as anti-oxidant and anti-arteriosclerotic possibility [13,14]. Several studies have shown that millet, as a kind of functional food, is playing an unignoring role in maintaining homeostasis of blood glucose [15,16], delaying gastric emptying [17,18], and enhancing immune competence. Clinical studies by Dewey and Ganmaa et al. have shown that dietary intervention can promote good absorption of nutrients and prevent or improve malnutrition [19,20,21]. However, how to effectively develop undernutrition-therapeutic diet with millet to remains to be further studied. This study aims to provide a functional or therapeutic nutritional product formula with millet, and to verify the improvement of intestinal, metabolic and immune functions in undernourished piglets.

## Materials and Methods

### Animals experimental design and procedures

Thirty-six female Durox × Danish Landrace × Yorkshire crossbred pigs (Shenzhen Agriculture and Animal Husbandry Co., Ltd.) were bought at an initial mean body weight of 3∼4 kg. Pigs were weaned at the age of 4 weeks and fed *ad labitum* with adequate formula for 1-week acclimatization. Then 28 pigs were switched from the formulated diet to a pure maize diet (Maize, n=28), while the other 10 pigs remained on the optimal formulated diet (Ref, n=8). After 4 weeks of respective feeding, the undernourished pigs were randomized into 3 groups based on the average body weight. In the following 3 weeks, malnourished pigs received CSB+ (CSB+, n = 10), MSB+ (MSB+, n = 10), maize (Maize, n = 8) diet accordingly while Ref continued on the optimal formulated diet as positive control. All the groups had *ad libitum* access to feed and water during the study period. Nutritional composition of diet was presented in Table 1. After refeeding, all the piglets were sacrificed for sample collection. The whole animal experiment was permitted and inspected by the BGI bioethics and biosafety board. CSB+ was produced completely according to the diet composition provided by the United States Agency for International Development (USAID), as well as MSB+ with slight difference because of the nature difference between corn and millet and the Standards for the Use of Food Nutrition Fortifiers (GB 14880).

**Table 1.**
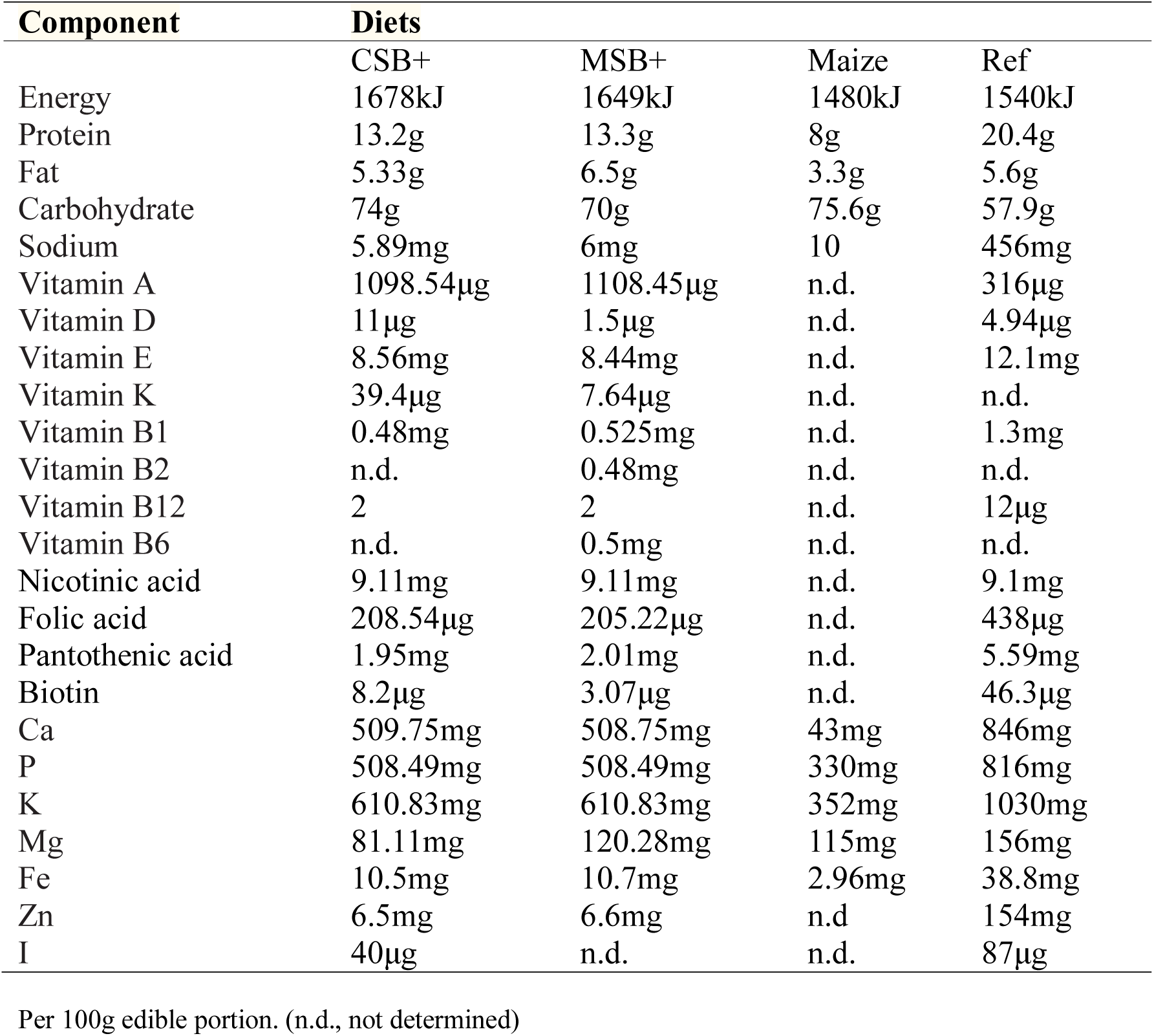
diets compositions.

### Growth and malnutrition estimation

Growth measurements, including body weight and crown-rump length (CRL), were collected weekly. These anthropometric measurements were used to assess the degree of malnutrition as described in details below. Based on the lengths of the undernourished pigs, theoretical weights were calculated (i.e. Weight in kg (theoretical) = e^3.77^ kg/m * [CRL length in meters] ^2.401^) and an estimate of the degree of wasting (i.e. Degree of wasting = (weight (observed)/weight (theoretical)) * 100%); The degrees of underweight and stunting were calculated as percentages of the means of the age-matched reference pigs: (i.e. Degree of underweight = (weight of the undernourished pigs/mean weight of the reference pigs) * 100%); Degree of stunting = (length of the undernourished pigs/mean length of the reference pigs) * 100%) [22]. Degrees of wasting, underweight and stunting were rated as severe, moderate or mild-normal in Table 2.

**Table 2.**
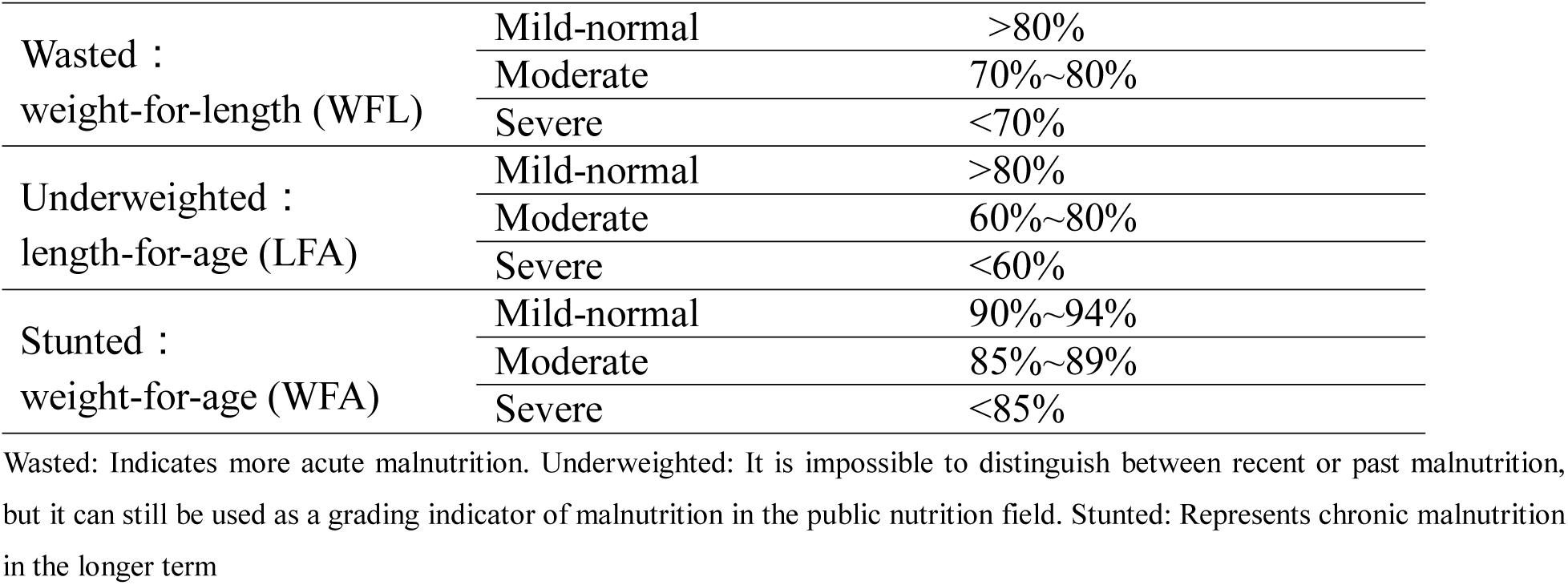
Assessment of degree of wasting, underweight and stunting.

|  |  |  |
| --- | --- | --- |
| Wasted :<br>weight-for-length (WFL) | Mild-normal | >80% |
|  | Moderate | 70%~80% |
|  | Severe | <70% |
| Underweighted :<br>length-for-age (LFA) | Mild-normal | >80% |
|  | Moderate | 60%~80% |
|  | Severe | <60% |
| Stunted :<br>weight-for-age (WFA) | Mild-normal | 90%~94% |
|  | Moderate | 85%~89% |
|  | Severe | <85% |

Wasted: Indicates more acute malnutrition. Underweighted: It is impossible to distinguish between recent or past malnutrition, but it can still be used as a grading indicator of malnutrition in the public nutrition field. Stunted: Represents chronic malnutrition in the longer term

### Blood sampling, tissue collection and histology assessment

For biochemical and systemic immune analysis, blood was collected by venipuncture of jugular venous from anaesthetized pigs just prior to euthanasia. For each sample, blood was transferred to a condensation and a heparin sodium tube respectively. Condensation tubes were centrifuged at 3500rpm for 15 min at 4°C and supernatant (serum) was harvested and stored at -80°C for blood biochemistry analysis. Similarly, heparin sodium tubes were centrifuged at 1600g for 10 min at 4°C and supernatant (plasma) was harvested for systemic immune analysis.

The heart, liver, kidneys, spleen and lungs were removed from the pigs and weighed immediately after euthanasia. Distal colons were fixed separately in 5% paraformaldehyde, and embedded with paraffin wax for histology assessment. The paraffin-embedded tissues were cut on a microtome (Leica RM2235, Germany) into 5μm sections and stained with hematoxylin and eosin. The whole tissue film was observed under the light microscope (OLMPUS CX22, Japan) and the histopathological changes were photographed by the microscopic imaging system (Leica DM1000, Germany). After the distal colon sections were stained with Alcian blue and Periodic acid-Shiff (AB-PAS), the ability of goblet cells to secrete mucus was evaluated by measuring the area of acidic mucin, neutral mucin and mucous layer and quantificating the relative areas of acidic mucin and neutral mucin to the total area of mucous layer.

### Biochemistry and systemic immune analysis

For blood biochemistry analysis, the levels of Albumin, total protein, urea, glucose, triglyceride, total cholesterol, low-density protein cholesterol, creatinine, C-reactive protein and total bile acid (all purchased from Maccura, Sichuan, except total bile acid from KHB, Shanghai) in serum were measured using an Automatic Analyzer 3100 (HITACHI, Japan). For systemic immune analysis, the levels of LPS, leptin (KENUODIBIO, SU-B50226; SU-B50089) and inflammatory factors (IL-1β, TNF-α, and IL-6) were determined using ELISA kits (AMEKO, AE90731Po; AE90301Po; AE90247Po) according to the manufacture’s instruction.

### Short-chain fatty acid (SCFA) determination by Gas chromatography-mass spectrometry (GC-MS)

Colon contents samples were prepared for GC-MS analyses as previously described with slight modification [23]. For colon contents, the extraction procedures were performed at 4 °C to protect the volatile SCFAs. In brief, 200g samples were mixed with 2mL of 5μg/ml ethyl acetate containing 2-Ethylbutyric acid as an internal standard. 200μL of 1M sodium chloride solution saturated with hydrochloric acid was added then. The mixed solution was ultrasound for 1h and then centrifuged for 10 min at 10000rpm at 4°C. MgSO_4_ was added to supernatant (20 times dilution) and then centrifuged (18000rpm, 10 min, 4°C). Collected supernatant 72μL and 18μL of MTBSTFA (Aladdin, Shanghai, China) was added. The solution was heated for 20 min at 80°C for derivatization. Then, the samples were cooled and transferred for GC-MS analyses.

GC-MS analyses were performed according to the method described by with several modifications [24]. The SCFA analysis was carried out using a TSQ 9000 GC-MS/MS. A nonpolar DB-5MS capillary column (30m × 0.25mm × 0.25μm, J&W Scientific, Folsom, CA, USA) was used for chromatographic separation. Helium (1.0ml/min) was used as the carrier gas. The chromatographic stepwise thermal conditions were as follows: 60°C for 2 min, 6°C/ min until 90°C for 2 min, 6°C/min until 120°C and 50°C/min until 280°C for 3 min. The mass spectrometer was set to scan mode from m/z 30–300 and in selected ion monitoring mode at m/z of 131 for propionic acid, m/z 145 for butyric acid and isobutyric acid, m/z 159 for isovaleric acid and valeric acid, and m/z 173 for 2-Ethylbutyric acid (internal standard).

### 16S rDNA Microbiome sequencing

Total cellular DNA was extracted with the E.Z.N.A. Stool DNA Kit (Omega) according to the company instruction. The bacterial hypervariable V3-V4 region of 16s rRNA was chosen for MiSeq (Illumina, CA USA) paired-end 300bp amplicon analysis using primer: 341_F: 5’-CCTACGGGNGGCWGCAG-3’ and 802_R: 5’-TACNVGGGTATCTAATCC-3’. The library preparation followed the method published previously [25].

### Statistics

All continuous data (except microbiome data) was analyzed in R (version 3.6.0) with respective packages. The biochemical and systemic immune data was analyzed by annova and multiple pairwise t-tests were adjusted by Turkey procedure in compareGroups (version 4.0) [26]. The intestine inflammation index and SCFA data were analyzed by Wilcoxon test for pairwise tests and the p values were adjusted by Benjamini-Hochberg procedure for each independent experiment, where R package rstatix (version 0.1.0) was implement to complete analysis [27]. Zeros in SCFAs data were replaced by the lowest value in all the samples. Data were shown as mean ± SD and mean ± SEM accordingly. Adjusted P values below 0.05 were regarded as statistically different.

For microbiome analysis, the raw sequencing reads were merged and trimmed, which followed by removing chimera and constructing zero-radius Operational Taxonomic Units (zOTUs) with UNOISE implemented in Vsearch (v2.6.0) [28, 29]. The green genes (13.8) 16s rRNA gene database was used as reference for annotation. Qiime2 (2018.11) combined with R packages(ggplot2, vegan) was used to process the forward analysis [30, 31, 32]. Rare zOTUs with frequency below 0.1% of the minimal sample depth were removed and filtered zOTU table was rarified to adequate sample depth (47000 count) for alpha and beta diversity calculation based on rarefaction curve. Principal coordinate analysis (PCoA) was conducted on unweighted and weighted Unifrac distance and permanova was performed to detect statistical difference between groups and p values were adjusted after pairwise tests. Specific taxa comparison among groups was analyzed by ANCOM and default setting in qiime2 was used to test statistically significant difference. For the different taxon found with ANCOM, Wilcoxon tests were conducted among the pairwise groups and the p values were adjusted with Benjamini-Hochberg procedure.

Rarified OTU table was used to predict functional gene with PICRUSt2 [33,34]. KO table was rarefied to the lowest sample depth and visualized by PCoA plot based on binary Jaccard and Bray Curtis distance. The KEGG Orthology (KO) table was downloaded from KEGG website (March, 2019) for annotation. Only the predicted KOs annotated in the KO table were remained to calculate the pathway abundance. The significantly abundant pathways were identified by LefSe with a strict threshold of linear discriminant analysis (LDA) score above 3 [35].

Rarefied OTU table was collapsed at the genera level and rare genera were removed according to a pre-set cut-off (mean relative abundance > 0.1% and appearance in more than 30% samples). Zeros were regarded as NA. Pearson correlation analysis implemented in Rhea was conducted between relative abundance of genera and the phenotype data after centered log-ratio data transformation. Correlation matrix was visualized with R package corrplot and correlation pairs with adjusted P value < 0.05 and absolute value of coefficient > 0.5 were plotted in scatter plot with linear regression using the default setting of Rhea [36, 37].

## Results

### Phenotype data analysis and SCFA quantification with GC-MS

After 3-week nutritional supplement, compared with Ref, CSB+, MSB+ and Maize fed piglets still remained low level of body weight and CR length, most were presenting severe wasted, underweighted and stunted (P < 0.01, Supplementary Table. 1). However, the malnutrition index didn’t show distinct difference among the undernutrition pig groups except for a lower weight-for-age index of MSB+ relative to Maize (Supplementary Figure. 1). Biochemical data showed significant differences among these experiment groups except for triglyceride and total bile acid (P < 0.05, Supplementary Table. 1). Compared with Maize, both CSB+ and MSB+ improved total protein and globulin in the serum (P<0.05, Table. 1) and globulin of MSB+ pigs remained at a similar level relative to Ref (P=0.70 and 0.51 respectively). Besides, urea was increased in CSB+ and MSB+ group due to high-protein feeding switch but showed no significant difference relative to Ref (P=0.06 and 0.15 respectively, Supplementary Table. 1). MSB+ had more a similar level of cholesterol with Ref in contrast to CSB+ (P=0.5 and 0.06 respectively, Supplementary Table. 1). Systemic immune dysbiosis existed in undernourished pigs with up-regulated IL-1beta and IL-6 relative to normally-grown pigs. MSB+ and CSB+ decreased IL-1beta and IL-6 but with no distinct different relative to Ref and Maize on account of wide range of data distribution. Meanwhile, MSB+ and CSB+ didn’t show significant difference in reverting the inflammation-dominant status (Supplementary Table. 1).

The pathological severity were evaluated according to histological characteristics of epithelial cells, mucosal and submucosal structures in the colon. Mild, moderate and severe pathological change was defined as shown in Table 3). In the maize group, 7 of 8 piglets showed pathological changes in the colon, and of which 5 presents mild, 1 was moderate and 1 was severe. However, pathological changes in 6 of 8 piglets in Ref were also observed, 4 had mild pathological changes and the rest presented moderate. In the refeeding groups, pathological changes were found in 8 piglets out of 10 in CSB+ and 6 out of 9 in MSB+ group. 7 were defined as mild and 1 as moderate in CSB+ group while 4 were mild, 1 moderate and 1 severe in MSB+ group (Table 4).

**Table 3.**
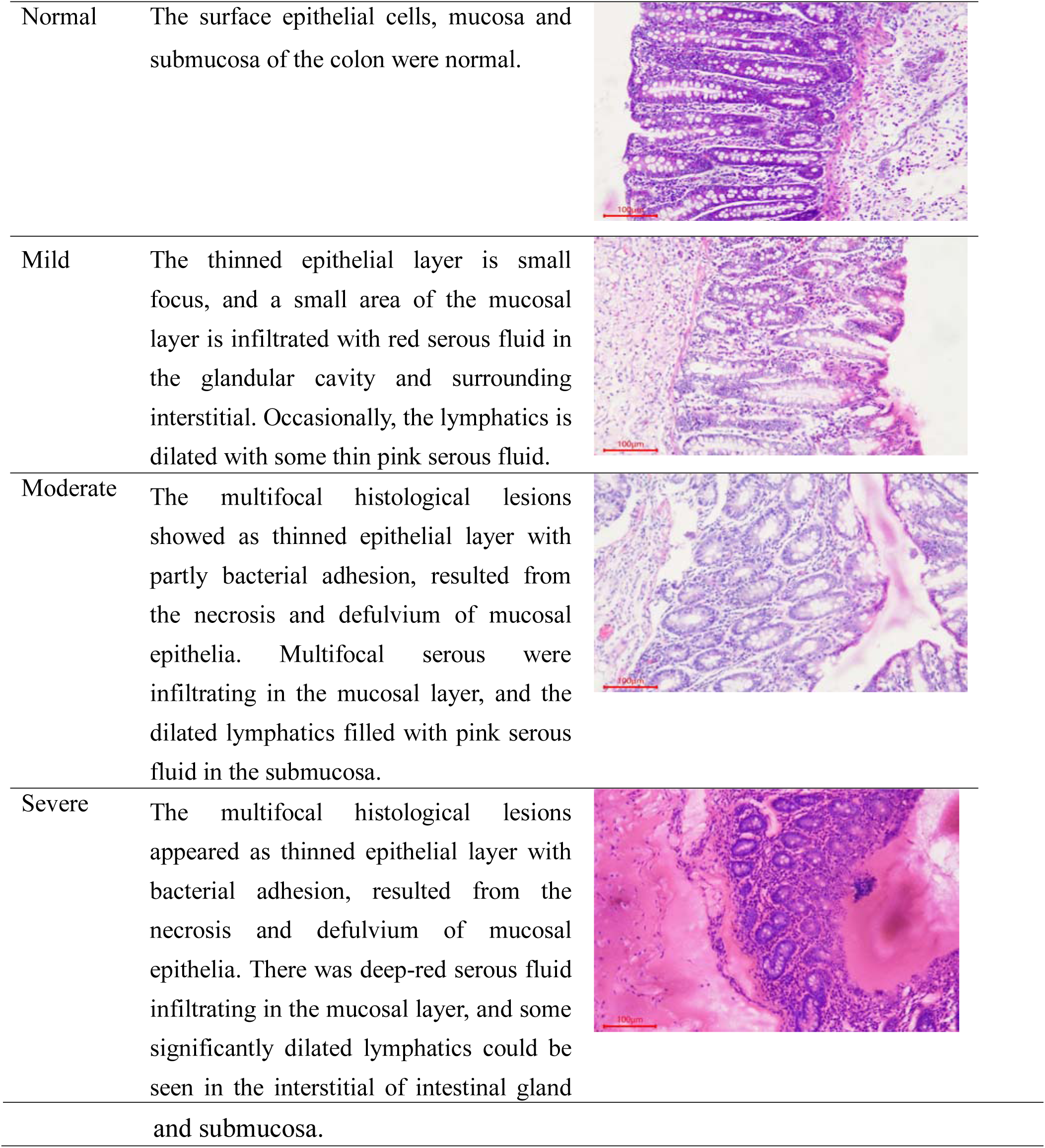
Histopathological scoring criteria of distal colon determined by evaluation of whole film.

**Table 4.** Histopathological levels of distal colon in respective experiment groups.

| Level | Maize | Ref | CSB+ | MSB+ |
| --- | --- | --- | --- | --- |
| Total | 8 | 8 | 10 | 9 |
| Normal | 1 | 2 | 2 | 3 |
| Mild | 5 | 4 | 7 | 4 |
| Moderate | 1 | 2 | 1 | 1 |
| Severe | 1 | 0 | 0 | 1 |

One missing sample from MSB+ for histological assessment

Meanwhile malnourished pigs had relatively thinner intestine wall compared with Ref but no significant difference was found except CSB+ (P<0.05, Supplementary Figure. 2). The density of both neutral and acidic goblet cells showed no significant difference among groups (Supplementary Figure. 3). But after re-feeding, MSB+ increased the neutral mucin-containing goblet cells rather than the acidic ones. Short-chain fatty acids in the colon of malnourished pigs were quantified by GC-MS (Figure. 1). Nutritional supplement increased the abundance of propionic acid, which was significantly up-regulated in the MSB+ group relative to Maize. Except that, both CSB+ and MSB+ showed no distinct difference in contrast with Maize.

**Fig 1.**
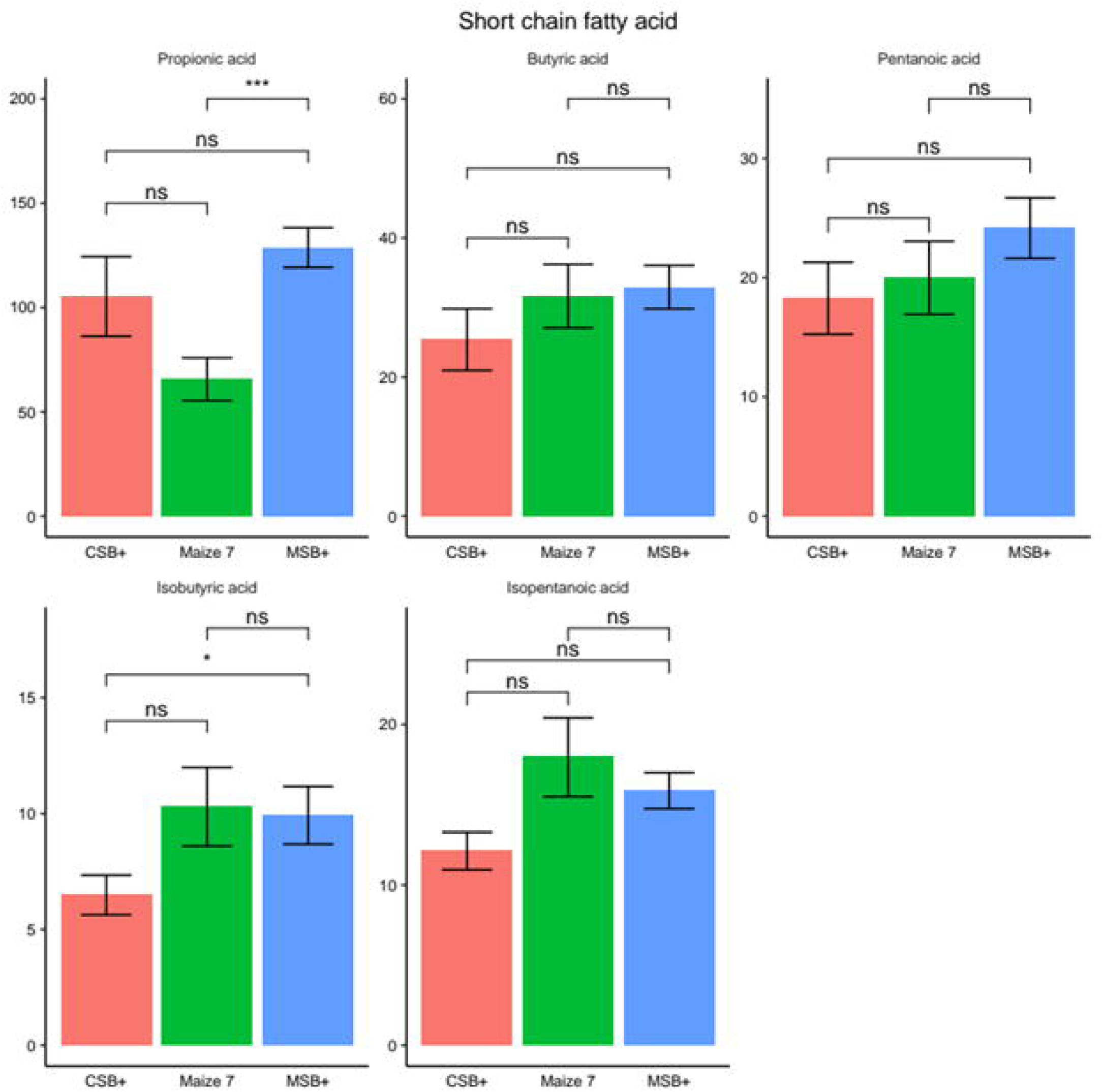
SCFA level in colon content determined by GC-MS (D). Data in the bar plot was shown by mean value together with SEM error bar. The Labels of ns, *, ** represents adjusted P > 0.05, < 0.05, < 0.01 respectively.

### Microbial structure profiling determined by 16s amplicon sequencing

Conlon microbial structure was determined by 16s amplicon sequencing. All the sequenced samples were rarefied to 47000 for the alpha and beta diversity comparison based on the rarefaction curve (Supplementary Figure. 4). Total observed zOTU number and Heip’s evenness were calculated to determine the ecosystem difference of diversity and evenness (Figure. 2A). Compared with Ref, Maize had less diverse and unbalanced microbial environment (P < 0.05 and 0.01, respectively). CSB+ and MSB+ feeding increased the malnourished pigs’ microbiota diversity (P < 0.01, 0.001, respectively). Moreover, MSB+ even had more observed zOTUs but remained similar level of evenness relative to Ref (P < 0.05 and > 0.05, respectively). Unweighted and weighted Unifrac distance were calculated to find that huge difference existed between the low-weight pigs and normal ones (Figure 2B and 2C). We conducted pairwise permanova tests on unweighted Unifrac distance and found all 4 groups were significantly different to each other (Adjusted P < 0.001). Nevertheless, CSB+ and Maize were more similar when giving weights to bacterial abundance (Adjusted P > 0.30, Figure. 2C). What’s more, different plant-based feeding led to a varied microbial abundance at phylum level (Figure. 2D). Firmicutes, Bacteriodetes and Proteobacteria were three main phyla found in the pigs, where no significant differences were found between MSB+ and Ref. In contrast to these two group, CSB+ however shared much more similar pattern with Maize (P > 0.05) which Proteobacteria overtook Bacteroidetes and Firmicutes. CSB+ and Maize had a relatively lower level of Firmicutes and Bacteriodetes, but the ratio in between didn’t show a distinct different among the four groups (Figure. 2E).

**Fig 2.**
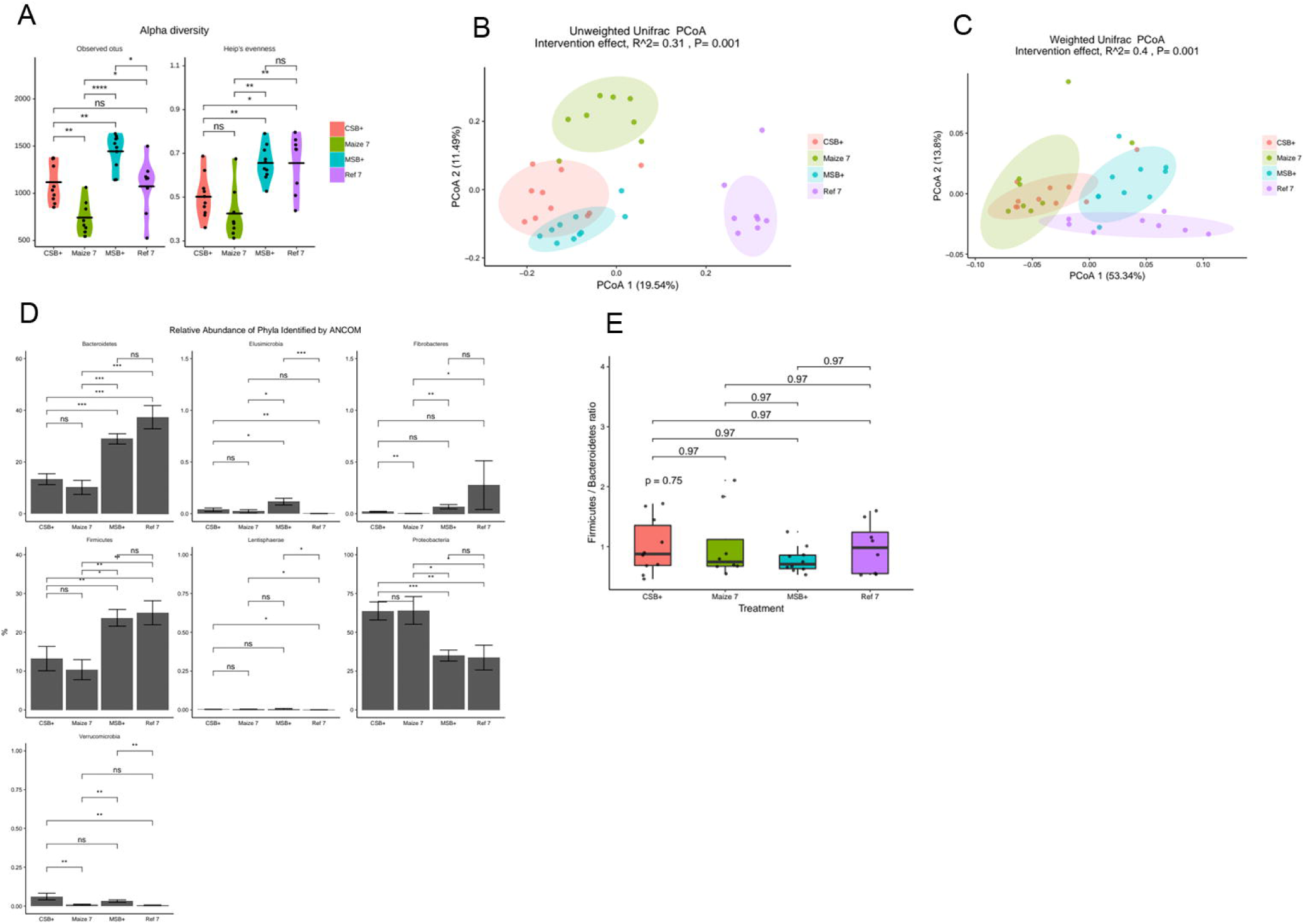
Gut microbiota determined by the 16s rRNA gene amplicon sequencing. Observed zOTU numbers and Heip’s evenness (A). PCoA plot of unweighted (B) and weighted (C) Unifrac distance metrics based on rarefied zOTU table, 80% confidence level at which to draw a respective ellipse following multivariate t-distribution. Relative abundance of different phyla identified by ANCOM (D). Relative abundance ratio between Firmicutes and Bacteriodetes (E). For CSB+ and MSB+, n=10; For Maize and Ref, n=8. Data in the bar plot was shown by mean value together with SEM error bar. The Labels of ns, *, **, ***, **** represent adjusted P > 0.05, < 0.05, < 0.01, < 0.005, < 0.001 respectively.

### Pearson’s correlation analysis between phenotype and gut microbiota

Rarefied zOTU table was collapsed at genus level with 103 annotated and 58 unambiguously identified to upper levels. *Campylobacter, Prevotella, Flexispira* and *Helicobacter* were the dominant genera in those immature pigs, taking up more than half of the microbial community. For CSB+ and Maize, *Campylobacter* and *Flexispira spp*. accounted for the high abundance of Proteobacteria while decreased levels of these in MSB+ and Ref were replaced by growing number of *Helicobacter* and other bacteria like *Provotella* from Firmicutes and Bacteriodetes (Figure. 3A). All the genera with relative abundance above 0.1% were kept for Pearson’s correlation analysis with phenotype after both were transformed with log-ratio. Body weight, CR length, total protein, albumin and creatinine were closely related to microbiota relative to other phenotype data. *Lactobacillus, Lachnospira, Prevotella, Faecalibacterium* were positively related with Pearson’s coefficient > 0.5 to the body weight, CR length, total protein except *Lactobacillus* and total protein (Pearson’s coefficient =0.38). Moreover, *Faecalibacterium* and *Lachnospira* were determined by ANCOM as significantly different genera among 4 groups (Figure. 3B). *Faecalibacterium* of MSB+ remained at a similar level with CSB+ and Maize (P > 0.05) while *Lachnospira* significantly increased (P < 0.05 and 0.01, respectively) after MSB+ feeding. What’s more, these two genera were highly correlated with malnutrition-associated phenotype with a significant adjusted P < 0.05 (Figure. 3C).

**Fig 3.**
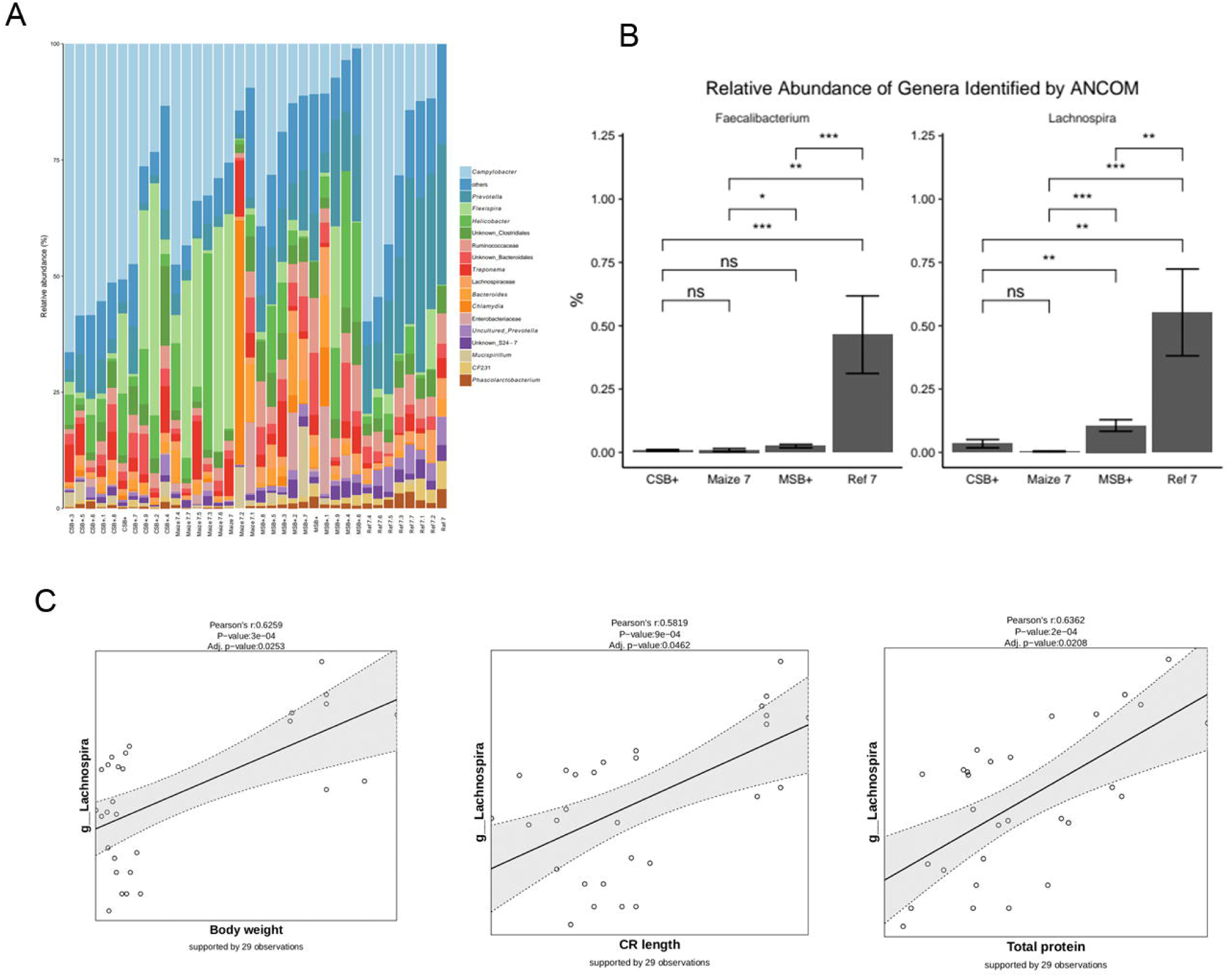
Data integration between microbiome data and phenotype. Relative abundance of prokaryotic gut microbiota members at genus level (A) and *Faecalibacterium* and *Lachnospira spp*. abundance in the gut content as determined by ANCOM (B). Scatter plot of the linear regression of the significant pairs about *Lachnospira spp*. and phenotype (C). For CSB+ and MSB+, n=10; For Maize and Ref, n=8. Data in the bar plot was shown by mean value together with SEM error bar. The Labels of ns, *, ** represents adjusted P > 0.05, < 0.05, < 0.01 respectively.

### Functional structure profiling with PICRUst2

Not only did microbial structure shift differently, the functional gene enriched pattern also drifted to varied direction (Figure. 4A and 4B). PICRUst2 was used to predict function KOs based on 16s copies and 6332 KOs were predicted with the minimum frequency of 32711943. All the KOs were rarefied to 32000000 to calculated the binary Jaccard and Bray Curtis distance metrics. Different plant-based feeding influenced the bacterial metabolism process. CSB+, MSB+ and Maize were not consistent from normal pigs (P < 0.01) when only considering whether one KO appear or not (Figure. 4A). However, we found MSB+ and Ref were more similar when using the abundance-weighted Bray Curtis distance (P = 0.07), which indicated some enriched KOs had similar abundance in the two groups (Figure. 4B). No distinct difference was detected between CSB+ and Maize using both binary Jaccard (P = 0.17) and Bray Curtis (P = 0.36) distance metrics.

**Fig 4.**
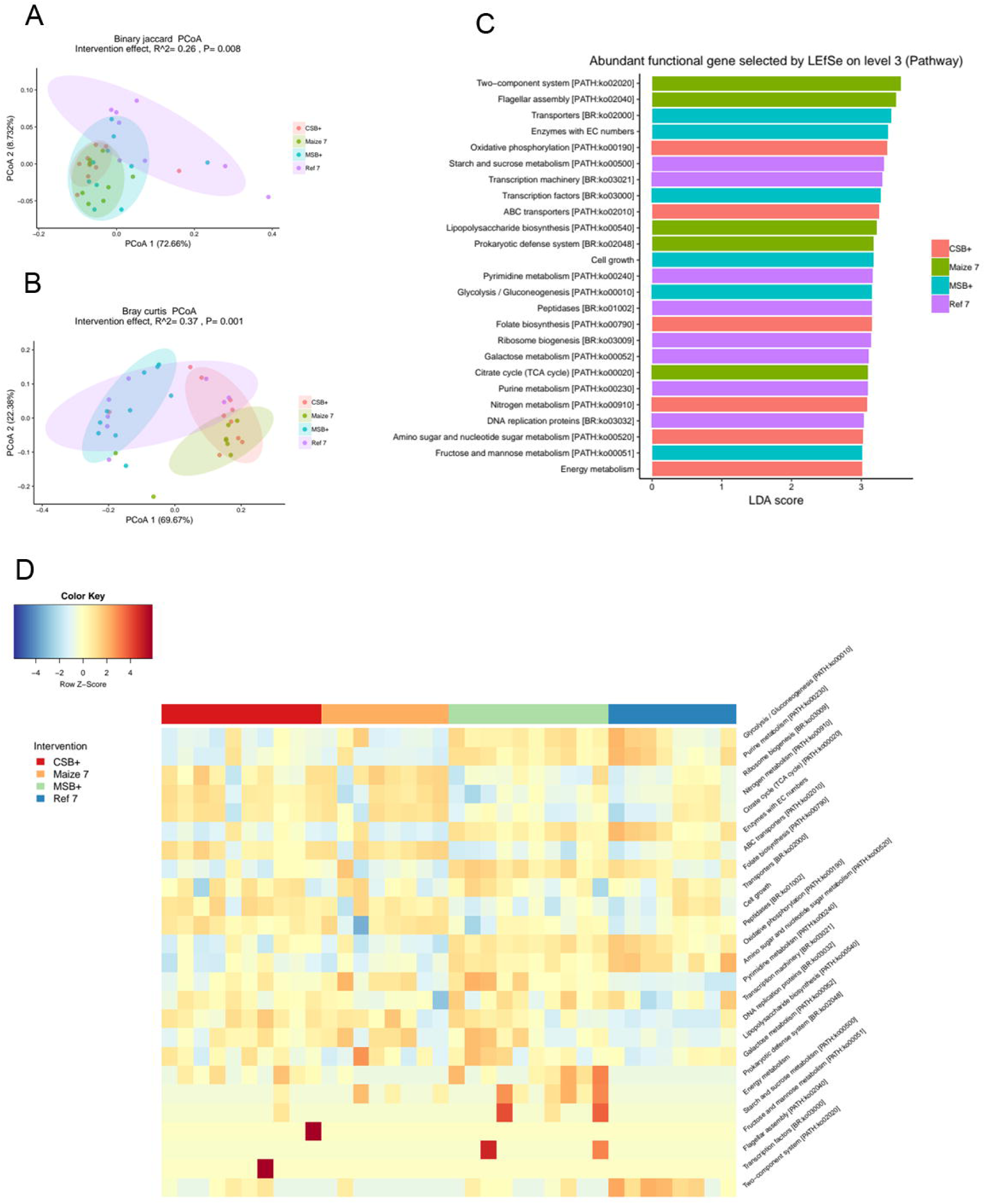
Functional gene prediction of the gut microbiota by PICRUSt2. PCoA plot of binary Jaccard (A) and Bray Curtis (B) distance metrics based rarefied KO table, 80% confidence level at which to draw a respective ellipse following multivariate t-distribution. LDA scores (C) and relative abundance (D) of the enriched pathways identified by LefSe. Relative abundance was visualized in heatmap after Z-score normalization. For CSB+ and MSB+, n=10; For Maize and Ref, n=8.

Bacterial metabolism pattern of malnourished pigs shifted to Ref after MSB+ feeding. In order to identify the difference of pathway enrichment, all the KOs were collapsed to 267 pathways and 24 respective pathways were identified with LDA score > 3 (Figure. 4C) and visualized in heatmap after Z-score normalization. MSB+ resembled the metabolic enrichment of Ref in contrast to CSB+ and Maize (Figure. 4D). Purine metabolism, glycolysis/gluconeogenesis, folate biosynthesis, oxidative phosphorylation and other bacterial metabolic pathways were enriched in Both MSB+ and Ref, compared to a relatively low level in CSB+ and Maize. Reversely, nitrogen metabolism, citrate cycle, and ABC transporters were mainly enriched in CSB+ and Maize but not in MSB+ and Maize.

## Discussion

### MSB+ and CSB+ didn’t revert the malnutrition status but readjusted the weak immune system in short-term intervention

Children are the most vulnerable to poor nutrition owing to poorly developed immune system and puberty lateness [38]. It has been regarded as complex to adopt appropriate therapeutic strategy for malnourished children. Malnourished children usually need to ingest enough calorie to sustain body development while they mostly face serious digestive problem or lack of appetite at the same time [39], which hampers restoring normal pace of body growth in later adolescence [40]. In this study, either millet-based or corn-based supplement didn’t make malnourished pigs recover to the same growth level relative to the well-fed in short-term intervention, led to unimproved physiological parameters including body weight and CR length which still indicated malnutrition among those re-fed piglets. Nonetheless, increase of total protein and globulin in the serum, as well as distinct increase of serum globulin was observed to reach the same level of well-fed pigs after nutritional intervention. Compared with Maize, relatively lower level of IL-1beta and IL-6 in CSB+ and MSB+ also indicated that nutritional intervention could adjusted the unbalanced immune system of the undernourished. Undernutrition was usually supposed to be followed by leaky gut and weaken immunity owing to insufficient food intake. However, we hardly observed any difference on pathological change between different groups, since even well-fed piglets also presented mild and moderate pathological changes. Moreover, both thickness of mucosa and density of goblet cells were only associated with the “size” of the piglets rather than the nutritional status. These data might be unsurprisingly explained why nutritional supplement from CSB+ and MSB+ seemingly invoked the weak immune system of malnourished pigs and adjusted the immune bias, which is also one important aspect in undernutrition therapy.

### Millet-based supplement restored well-fed like microbiota of the malnourished

Despite the physiological and immune assessment, gut microbiota has been listed as one important area in practical malnutrition intervention [12, 41]. In the study, CSB+ and MSB+ both significantly improved the diversity of microbiota while the evenness of ecosystem was also restored after MSB+ intervention. Compared with CSB+, MSB+ had more diverse and evenly-distributed commensal microbiome, that is to say, a steadier system. Moreover, MSB+ had more well-fed like microbial structure with higher level of Bacteriodetes and Firmicutes rather than Proteobacteria. Similarly, a decreased diversity and enrichment of potentially pathogenetic Proteobacteria were previously reported to appear in undernourished children’s microbiota [42,43].

Decreased abundance of *Campylobacter* and *Flexispira* explained the missing part from Proteobacteria in MSB+ and Ref, which was largely taken by *Helicobacter, Prevotella* and other bacteria from Firmicutes and Bacteriodetes. Besides those, the level of propionic acid in colon, relative to Maize, was found significantly high after MSB+ supplement, which also confirmed the commensal bacteria in an effective order of cooperation. However, considering CSB+ and Maize are both corn-based feeding, corn-based supplement like CSB+ might have some drawbacks to help the malnourished regain well-fed microbiota compared with MSB+.

### *Faecalibacterium and Lachnospira spp*. was highly correlated with malnutrition-associated phenotype

Pearson’s correlation analysis indicated microbiota was highly correlated to malnutrition-associated phenotype, especially body weight, CR length, total protein, albumin and creatinine. Potentially beneficial *Lactobacillus, Lachnospira, Faecalibacterium and Roseburia spp*. were all found positively correlated with these index (Pearson’s correlation > 0.5), supporting the microbiota dysbiosis of the malnourished. Previously, children with low levels of *Lachnospira, Veillonell, Faecalibacterium* and *Rothia* in their first 3 months were discovered to have higher risk for asthma and tend to receive more antibiotics than healthier children before they turned 1 year old [44]. These missing beneficial bacteria explained the weaken immunity of in their childhood. Possibly, the low abundance of potentially probiotics might be one significant cause of malnourished piglets’ weak immune resistance. Promisingly, MSB+ significantly increased the abundance of *Lachnospira spp*. relative to CSB+ and Maize, which was also previously discovered to play an important anti-inflammatory effect in weaner pigs fed with a high-fiber diet [45].

### MSB+ pigs shared similar functional gene enrichment in microbial community relative to Ref

Along with microbial structure, MSB+ pigs also showed a similar metabolic pattern relative to well-fed piglets. Purine metabolism, glycolysis/gluconeogenesis, folate biosynthesis and oxidative phosphorylation were enriched in both groups, referring to an effective microbial cooperation system. Sufficient intake of nutritionally-balanced food allows commensal microbes to complete necessary metabolism and biosynthesis, which nurtures the host to some extent. More and more researches indicated host’ nutrition source not only comes from food intake from outside environment, but commensal microbes also provide mirco-nutrients like folate through metabolism of undigested food residue. Glycolysis/gluconeogenesis and oxidative phosphorylation provide the needed energy for microbe to complete normal growth and routine activities [46, 47]. Unlike MSB+ and Ref, nitrogen metabolism, citrate cycle, and ABC transporters were main pathways abundant in CSB+ and Maize, indicating the metabolic direction of microbial community differs upon the source of intake. Bacterial ABC transporter has been regarded as important roles in nutrient uptake and drug resistance, but increasing evidences point out either direct or indirect role roles in the virulence of bacteria [48]. Enriched nitrogen metabolism and possibly frequent ABC transportation revealed a high nutritional demand of bacteria together with a relatively monotonous biological structure, which might be a microbiota-disorder like phenomenon with some overgrown potentially pathogenetic bacteria.

To avoid the variation caused by the diet composition difference, we tried to minimize the difference between MSB+ and CSB+. However, the natural property of millet still increased the vitamin B2, B6 and magnesium in MSB+, and decreased vitamin D, vitamin K, biotin and iodonium because it is not allowed to add in the food as fortifiers according to the Standards for the Use of Food Nutrition Fortifiers (GB 14880). Therefore, the beneficial effects might count for the diet components of different amount. Moreover, as provide by many studies that millet is rich in bioactive components, such as antioxidants, antimicrobial components, which may also lead to better recovery to undernourished piglets. Further studies would be carried out to figure out the main effective components [49].

## Conclusion

In conclusion, this study discovered corn- and millet-based nutritional supplements didn’t revert the undernourished state of pigs in 3 weeks but positively modulated the immune system. Moreover, undernourished pigs shared a phylogenetically and functionally similar structure of microbiota with millet-based supplement. The study discovered the promising potentially of millet as material source in modulating the gut microbiota disorder of the malnourished. It would be relevant to validate our finding in other animal models or human cohort. Additionally, millet is drought-resistant and widely grown in arid and semiarid areas of Asia and Africa, where millions of children suffered from undernutrition problems. This study will attract more attention to further study on the practical use of the easy-to-get agriculture product in those areas.

