## Supplemental Table and Figure for "Millet-based supplement restored microbiota diversity of acute undernourished pigs"

Supplementary materials

Table 1. Serum biochemical and systemic immune index.

|  | **CSB+ N=10** | **Maize**  **N=8** | **MSB+ N=10** | **Ref**  **N=8** | **p.CSB+ vs Maize** | **p.MSB+ vs Maize** |
| --- | --- | --- | --- | --- | --- | --- |
| Body weight (kg) | 6.00±1.51 | 7.20±1.08 | 5.63±1.07 | 28.8±3.82 | 0.63 | 0.41 |
| CR length (cm) | 49.4±3.60 | 53.1±3.73 | 51.6±4.86 | 71.2±1.75 | 0.19 | 0.86 |
| Total protein (g/L) | 53.1±4.75 | 44.6±5.05 | 53.1±5.50 | 69.7±5.87 | 0.01 | 0.01 |
| Albumin (g/L) | 14.4±1.90 | 11.9±1.82 | 15.0±1.74 | 28.9±4.02 | 0.18 | 0.06 |
| Globulin (g/L) | 38.7±3.47 | 32.7±3.44 | 38.1±4.76 | 40.8±4.32 | 0.02 | 0.04 |
| Albumin/Globulin | 0.37±0.04 | 0.37±0.03 | 0.40±0.06 | 0.72±0.12 | 1.00 | 0.77 |
| Glucose (mmol/L) | 5.21±1.84 | 5.99±0.79 | 7.11±1.71 | 6.58±0.97 | 0.68 | 0.38 |
| Urea (mmol/L) | 6.89±2.53 | 5.47±1.90 | 6.55±1.00 | 4.77±0.60 | 0.32 | 0.56 |
| Creatinine (mmol/L) | 50.4±16.5 | 60.7±15.6 | 50.7±12.4 | 93.6±5.70 | 0.39 | 0.42 |
| Cholesterol (mmol/L) | 2.40±0.46 | 2.37±0.31 | 2.69±0.36 | 3.02±0.79 | 1.00 | 0.55 |
| Triglyceride (mmol/L) | 0.61±0.34 | 0.48±0.15 | 0.53±0.15 | 0.66±0.31 | 0.70 | 0.98 |
| Low-density protein (mmol/L) | 0.98±0.20 | 1.06±0.21 | 1.10±0.27 | 1.54±0.57 | 0.96 | 0.99 |
| C-reaction protein (mmol/L) | 8.42±4.29 | 4.69±2.82 | 3.73±1.65 | 5.25±3.31 | 0.08 | 0.92 |
| Total bile acid (μmol/L) | 24.6±9.70 | 33.4±24.3 | 44.3±23.2 | 28.5±11.2 | 0.74 | 0.60 |
| IL-1beta (ng/L) | 66.9±63.2 | 143±84.1 | 87.1±46.1 | 48.2±58.8 | 0.07 | 0.26 |
| IL-6 (ng/L) | 7.57±7.71 | 10.8±5.18 | 8.95±5.67 | 1.78±1.73 | 0.64 | 0.90 |
| TNF-alpha (pg/mL) | 19.2±5.52 | 18.7±11.0 | 22.4±4.90 | 16.4±8.37 | 1.00 | 0.74 |
| Leptin (ng/mL) | 22.2±9.86 | 15.8±9.95 | 20.8±11.2 | 17.8±8.58 | 0.54 | 0.71 |
| Lipopolysaccharide (pg/mL) | 1032±587 | 669±465 | 925±626 | 722±373 | 0.49 | 0.74 |

Descriptive data are shown as mean ± SD.

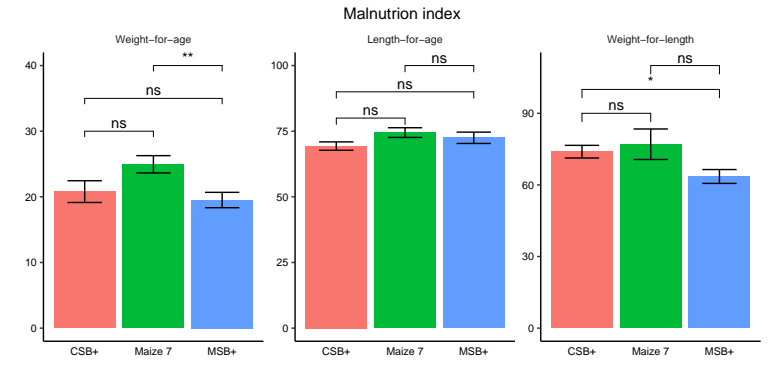

Figure. 1 Malnutrition index. The Labels of ns, *, ** represents adjusted P > 0.05, < 0.05, < 0.01 respectively.

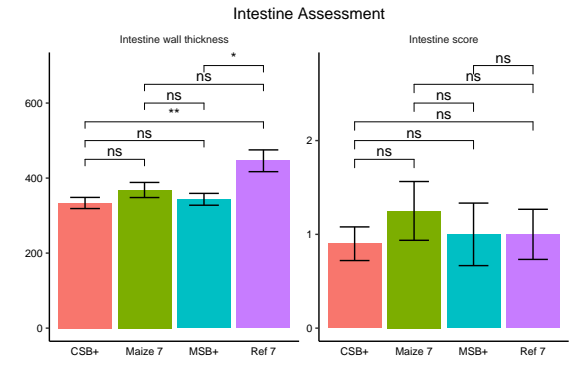

Figure. 2 Intestine wall thickness and intestine score. The Labels of ns, *, ** represents adjusted P > 0.05, < 0.05, < 0.01 respectively.

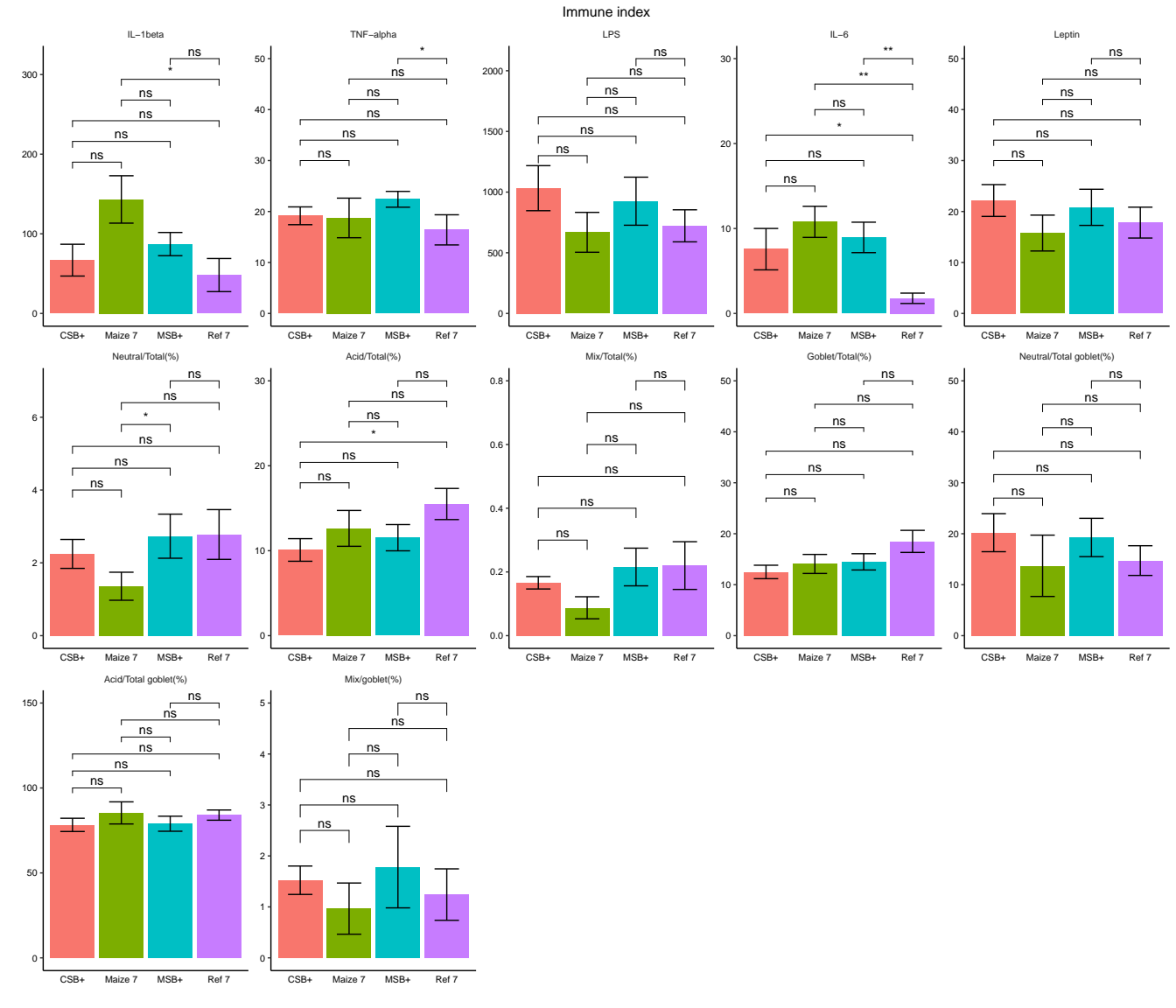

Figure. 3 Immune index. The Labels of ns, *, ** represents adjusted P > 0.05, < 0.05, < 0.01 respectively.

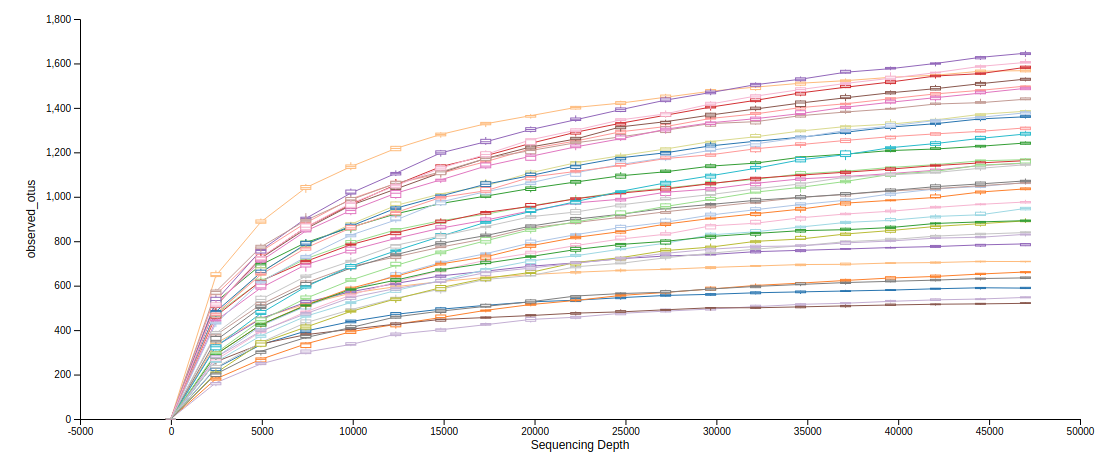

Figure. 4 Rarefaction curve
